# On the use of the experimentally determined enzyme inhibition constant as a measure of absolute binding affinity

**DOI:** 10.1101/144204

**Authors:** Fouad H. Darras, Yuan-Ping Pang

**Affiliations:** Computer-Aided Molecular Design Laboratory, Mayo Clinic, Rochester, MN 55905, USA

**Keywords:** dissociation constant, inhibition constant, affinity, nonspecific binding, nontarget site binding, adsorption

## Abstract

Defined as a state function representing an inhibitor’s absolute affinity for its target enzyme, the experimentally determined enzyme inhibition constant (*K*_i_) is widely used to rank order binding affinities of different inhibitors for a common enzyme or different enzymes for a common inhibitor and to benchmark computational approaches to predicting binding affinity. Herein, we report that adsorption of bis(7)-tacrine to the glass container surface increased its *K*_i_ against *Electrophorus electricus* acetylcholinesterase (*ee*AChE) to 3.2 ± 0.1 nM (n = 5) compared to 2.9 ± 0.4 pM (n = 5) that was determined using plastic containers with other assay conditions kept the same. We also report that, due to binding or “adsorption” of bis(7)-tacrine to the inactive *ee*AChE, the bis(7)-tacrine *K*_i_ increased from 2.9 ± 0.4 pM (n = 5) to 734 ± 70 pM (n = 5) as the specific *ee*AChE activity decreased from 342 U/mg to 26 U/mg while other assay conditions were kept the same. These results caution against using *K*_i_s to rank order binding potencies, define selectivity, or benchmark computational methods without knowing detailed assay conditions.

**Abbreviations:**

*K*_i_: enzyme inhibition constant
AChE: acetylcholinesterase
*ee*AChE: *Electrophorus electricus* AChE
ATCh: acetylthiocholine chloride
bis(7)-tacrine: 1,7-*N*-heptylene-bis-9,9'-amino-1,2,3,4-tetrahydro-acridinium dihydrochloride
DTNB: 5,5’-dithiobis(2-nitrobenzoic acid)
SEA: specific enzyme activity
tacrine: 9-amino-1,2,3,4-tetrahydroacridinium monohydrochloride.

## 1. Introduction

Enzyme inhibition constant (*K*_i_), also known as inhibitor dissociation constant, is an equilibrium constant of a reversible inhibitor for complexation with its target enzyme. Unless otherwise specified all inhibitors described hereafter are reversible inhibitors. *K*_i_ is associated with thermodynamic parameters in that Δ*G* = *RT*ln(*K*_i_), where Δ*G, R*, and *T* are the absolute binding free energy, the gas constant, and the absolute temperature, respectively [1]. Here *K*_i_ should not be confused with *K*_I_ of an irreversible inhibitor, which is the irreversible inhibitor concentration that causes a rate of inactivation equal to a half of pseudo-unimolecular inhibition rate constant. Nor should *K*_i_ be confused with *k*_i_ of an irreversible inhibitor that is a bimolecular inhibition rate constant [2–5]. Unlike the inhibitor concentration that causes 50% enzyme inhibition (*IC*_50_), *K*_i_ is a state function that is independent of the concentration of enzyme used to determine the *K*_i_. Therefore, *K*_i_ represents the absolute affinity of an inhibitor for its target enzyme, and one can theoretically use *K*_i_ to rank order binding affinities of different inhibitors for a common enzyme, define selectivity of an inhibitor for different enzymes, and benchmark in silico approaches to prediction of inhibitor binding affinities.

However, a cursory literature search showed a wide range of experimentally determined *K*_i_s for 9-amino-1,2,3,4-tetrahydroacridinium monohydrochloride (tacrine, a withdrawn Alzheimer’s drug [6]) against acetylcholinesterase (AChE) [7–13] from the same species using the same spectrophotometric Ellman assay [14] under the same assay conditions (temperature, pH, and ionic strength). For example, the *K*_i_ of tacrine was reported to be 20.2 ± 0.1 nM by one group and yet 340 ± 97 nM by another group for inhibiting *Electrophorus electricus* AChE (*ee*AChE) under the same Ellman assay conditions [15,16]. For another example, the *K*_i_ of tacrine was reported to be 36 ± 1 nM by one group and later 137 nM by the same group for inhibiting recombinant human AChE under the same Ellman assay conditions [17,18]. These results raised concerns on the use of the experimentally determined *K*_i_ as a measure of absolute binding affinity.

In this article we report our enzyme kinetics studies using a model system of *ee*AChE and its water-soluble inhibitors tacrine and 1,7-N-heptylene-bis-9,9'-amino-1,2,3,4-tetrahydro-acridinium dihydrochloride, an analog of tacrine known as bis(7)-tacrine [19], to evaluate the suitability of using the experimentally determined *K*_i_ as a measure of absolute binding affinity. The advantages of this model system are that AChE is a well-studied one-substrate enzyme and that preparation of the inhibitor solution for tacrine and bis(7)-tacrine does not require use of any co-solvent such as dimethyl sulfoxide, a mild oxidation reagent [20] that has an inhibitory effect on AChE [21].

## 2. Materials and methods

### 2.1. Materials

*ee*AChE was purchased from Sigma-Aldrich (St. Louis, MO; catalog number of C2888 with log numbers of SLBN0954V and SLBS4398 and specific enzyme activity of ≥1000 U/mg; catalog number of 3389 with log number of SLBL3186V and specific enzyme activity of 200–1000 U/mg). Acetylthiocholine chloride (ATCh), NaH_2_PO_4_, Na_2_HPO_4_, and Triton X-100 were purchased from ACROS (Morris Plains, NJ). 5,5’-Dithiobis(2-nitrobenzoic acid) (DTNB) and tacrine were ordered from Sigma-Aldrich (St. Louis, MO). Bis(7)-tacrine was synthesized according to a published scheme [19]. Inhibitor purity was confirmed by elemental analysis performed at NuMega (San Diego, CA). Tacrine: Anal. Calcd. for C_13_H_17_ClN_2_O: C, 61.78; H, 6.78; N, 11.08. Found: C, 61.57; H, 7.20; N, 11.17. Bis(7)-tacrine: Anal. Calcd. for C_32_H_44_Cl_2_N_4_O_2_: C, 65.41; H, 7.55; N, 9.53. Found: C, 65.81; H, 7.63; N, 9.34.

### 2.2. Specific enzyme activity and K_i_ determination

Briefly, to each of 40 wells in a flat-bottom, clear, 96-well plate was added at 26 °C sequentially 270 μL 50 mM sodium phosphate buffer (pH 8.0) with 0.1% (v/v) Triton X-100, 5 μL *ee*AChE solution (15.000, 7.5000, 5.000, 2.500, or 0.625 μg/mL), 5 μL of inhibitor solutions (for tacrine: 3.0 μM, 1.5 μM, and 0.6 μM for 0.625 μg/mL of *ee*AChE or 6.0 μM, 3.0 μM, and 1.5 μM for 15.000 μg/mL of *ee*AChE; for bis(7)-tacrine: 0.6 nM, 0.3 nM, and 0.15 nM for 0.625 μg/mL of *ee*AChE or 90 nM, 60 nM, and 30 nM for 15.000 μg/mL of *ee*AChE) or 5 μL of distilled water (for control and the specific enzyme activity determination), 10 μL 2.5 mM DTNB, and 10 μL ATCh solutions (15.000, 7.500, 3.750, 1.875, and 0.938 mM). The resulting solutions were left on the bench at 26 °C for equilibration for 2 minutes and then measured for ATCh hydrolysis rate (*v*) at a microplate reader temperature of 26 ± 2 °C. The specific enzyme activity (SEA) for *ee*AChE was calculated according to SEA = (*A*•*V*)/(*ε*•*L*•*T*•*W*_E_), where *A* was the UV absorption in optical density (OD) of the ATCh hydrolysis product (0.21−1.26 × 10^−3^ OD); V was the volume of the assay solution (300 μL); ε was molar absorptivity at 405 nm (13.3 L•cm^−1^•mol^−1^) [22]; *L* was the length of the light path of the flat-bottom, clear, 96-well plate (0.75 cm); *T* was the time over which the hydrolysis product was generated (10 minutes); *W*_E_ was the weight of *ee*AChE (10.4–250 pg); 1U is defined as converting 1 μmol of substrate to its product in a minute [23]. *K*_i_ was obtained from 1/*v*, 1/[ATCh], and [tacrine] or [bis(7)-tacrine] using Prism 4 with the Lineweaver-Burk plot [24] (see Supplementary information for details).

### 2.3. UV absorptions of inhibitor solutions that were prepared using glass or plastic vials

Briefly, to a single quartz cuvette that was washed with distilled water and dried by blowing N_2_ gas, 3.0 mL of a tacrine or bis(7)-tacrine solution of 30.0 μM, 20.0 μM, 15.0 μM, 10.0 μM, 7.5 μM, or 5.0 μM was added using a 1000-μL Pipetman P1000 pipette. The cuvette with the highest tacrine or bis(7)-tacrine concentration was first placed in the SpectraMax Plus 384 Absorbance Microplate Reader to scan for λ_max_ from 190 nm to 400 nm. The λ_max_s for tacrine and bis(7)-tacrine were found to be 242 nm and 244 nm, respectively. The UV absorption of an inhibitor solution, which was prepared using two 2.0-mL microcentrifuge tubes or a 7.4-mL glass vial, was then determined by the observed absorbance of an inhibitor solution with or without 0.4% (v/v) Polysorbate 20 subtracted by the observed absorbance of distilled water with or without Polysorbate 20, respectively. The UV absorbance of an inhibitor at each concentration shown in Figure 1 was an average of at least three measurements each of which used a freshly prepared inhibitor solution (see Supplementary information for details).

**Figure 1.**
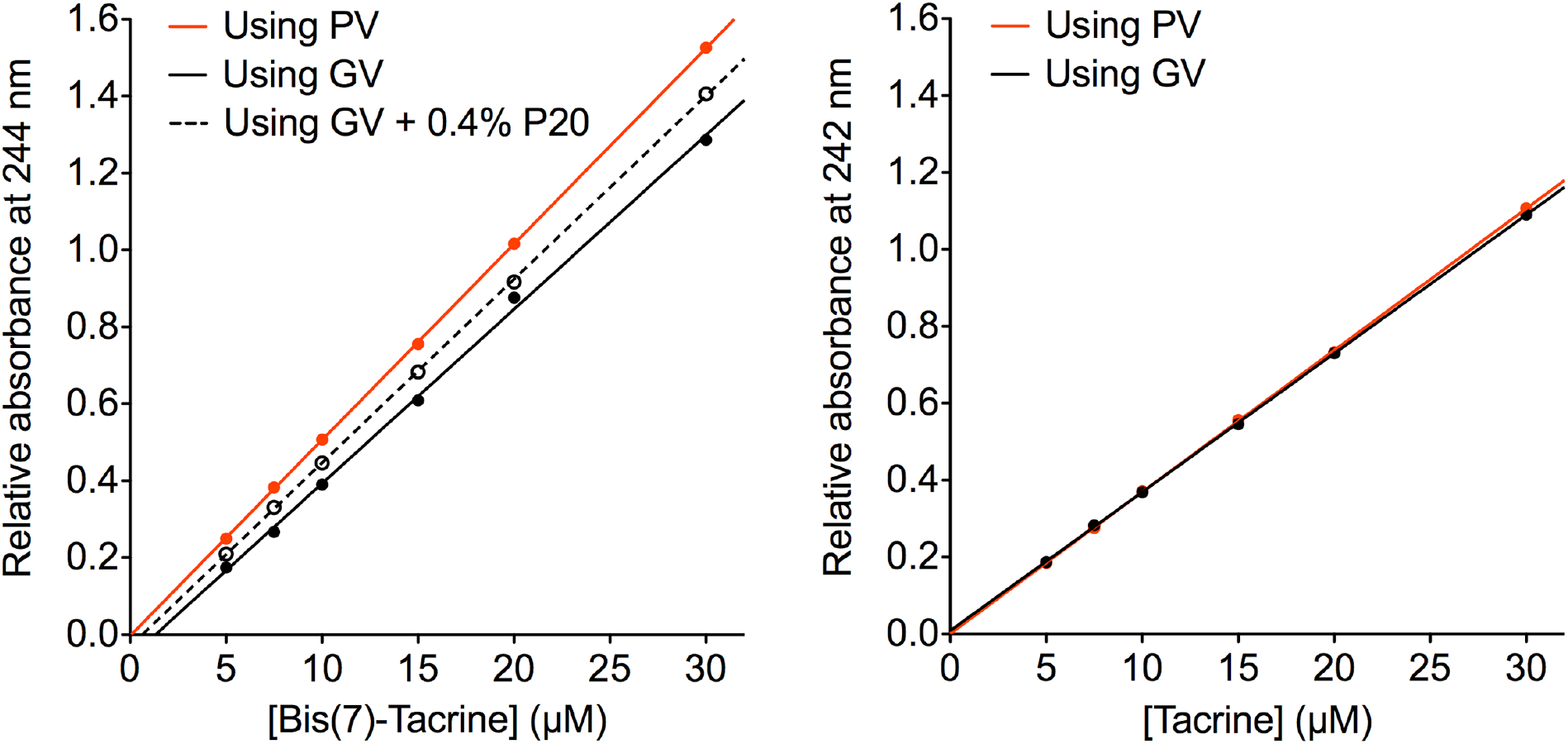
Relative UV absorptions of acetylcholinesterase inhibitor solutions that were prepared using glass or plastic vials with or without surfactant Polysorbate 20. Glass vials, plastic vials, and Polysorbate 20 are abbreviated as GV, PV, and P20, respectively. The relative absorbance was the absorbance of an inhibitor in distilled water with or without P20 subtracted by the absorbance of distilled water with or without P20, respectively. All UV absorptions were measured using a single quartz cuvette. The data for this figure are listed in Table S2.

## 3. Results and discussion

### 3.1. Container surface as nontarget binding site

To evaluate the suitability of using the experimentally determined *K*_i_ as a measure of absolute binding affinity, we first turned our attention to a seemingly trivial detail—the arbitrary use of either glass or plastic containers for stock solutions of AChE inhibitors in our enzyme inhibition studies. Adsorption of peptides or proteins to container surfaces and its effect on enzyme inactivation had been documented [3,25,26]. Additives leaching from laboratory plasticware had also been reported [27]. However, we did not find a report on adsorption of small-molecule inhibitors to container surfaces and its effect on *K*_i_. This led us to determine whether there was a difference in *K*_i_ for two inhibitor stock solutions that were prepared using a 7.4-mL general-purpose borosilicate glass threaded vial (the glass vial) and a widely-used 2.0-mL microcentrifuge tube (the plastic vial). Unexpectedly, we found the mean and standard error of *K*_i_ for bis(7)-tacrine against *ee*AChE to be 3.2 ± 0.1 nM (n = 5) or 2.9 ± 0.4 pM (n = 5) when the inhibition constant was determined using the glass or the plastic vials, respectively, while all other assay conditions were kept the same (Table 1). We also observed that the specific *ee*AChE activity resulting from short-exposure to the plastic vial (342 ± 10 U/mg; Table 1) was similar to the activity from the glass vial (334 ± 11 U/mg; Table 1). Rather than effects of possible additives leaching from the plastic vial, the 1000-fold difference in *K*_i_ indicated that substantially more adsorption of bis(7)-tacrine to the glass than plastic surface occurred during the inhibitor solution preparation process. This adsorption was confirmed by the differential UV absorptions of two bis(7)-tacrine solutions that were prepared using the glass and plastic vials (Fig. 1). It was further confirmed by the reduction of the difference in UV absorption that was caused by adding 0.4% (v/v) Polysorbate 20, a nonionic surfactant that was routinely used to prevent analytes from adsorption to the microfluidic system in Biacore-based surface plasmon resonance studies (Fig. 1). It is worth noting that complete desorption of bis(7)-tacrine is impossible because increasing the concentration of Polysorbate 20 reduces the water solubility of bis(7)-tacrine. Repeating the adsorption experiments using tacrine showed only slight differences in *K*_i_ and UV adsorption (Table 1 and Fig. 1). These results demonstrate that container surface can serve as a nontarget binding site for a test inhibitor during the inhibitor solution preparation process. The results also explain that the 1000-fold change for the *K*_i_ of bis(7)-tacrine against *ee*AChE was due to the adsorption-caused reduction of the actual inhibitor concentration that was available to *ee*AChE relative to the nominal inhibitor concentration.

**Table 1.** Effect of a stock solution container on *K*_i_

| Inhibitor | SEA<sup>a</sup> in U/mg (mean ± SE<sup>b</sup>) | | $K_i$<sup>c</sup> (mean ± SE<sup>b</sup>) | |
| --- | --- | --- | --- | --- |
|  | Glass | Plastic | Glass | Plastic |
| Bis(7)-tacrine | 334 ± 11 | 342 ± 10 | 3.2 ± 0.1 nM | 2.9 ± 0.4 pM |
| Tacrine | 122 ± 7 | 124 ± 7 | 35 ± 3 nM | 32 ± 1 nM |

<sup>a</sup>SEA: specific enzyme activity of *Electrophorus electricus* acetylcholinesterase that was determined using the data in Table S1 at pH = 8.0, 50 mM sodium phosphate buffer, temperature of 26 ± 2 °C, and acetylthiocholine concentration of 2.0 mM. <sup>b</sup>SE: standard error of five independent experiments. <sup>c</sup> $K_i$ was determined using the data in Table S1 and Fig. S1 employing glass or plastic vials for inhibitor stock solutions.

### 3.2. Inactive enzyme as nontarget binding site

In performing the study described above, we also observed a correlation for both tacrine and bis(7)-tacrine between the experimentally determined *K*_i_ and the amount of *ee*AChE required for the *K*_i_ determination assay (Table 2). In theory, *K*_i_ is independent of the amount of the enzyme used in the assay. However, the correlation indicated that in practice *K*_i_ depended on the amount of the enzyme used in the assay. The observed dependence of *K*_i_ on the amount of enzyme suggested that a test inhibitor might bind to not only the active enzyme but also the inactive enzyme caused by dilution denaturation and/or thermal inactivation during the process of the enzyme inhibition assay. For simplicity we do not consider herein the minor binding of the inhibitor to nonactive-site regions of the active enzyme and nontarget proteins that coexist with *ee*AChE. The binding to the inactive enzyme might consequently reduce the actual inhibitor concentration available to the active enzyme causing an overestimation of the experimentally determined *K*_i_ (*viz*., underestimation of the binding affinity). One way to confirm the inhibitor binding with the inactive enzyme was to confirm that the actual specific enzyme activity [23], a measure of the percentage of the active enzyme, was inversely proportional to *K*_i_. So we tested tacrine and bis(7)-tacrine using different batches of *ee*AChE—each of which had a unique specific enzyme activity that was determined at the time when the assay was performed—under the following three specific conditions. First, because loss of enzyme activity can occur while measuring reactions for extended periods [3] and each *K*_i_ determination took about 30 minutes to complete, we performed five independent *K*_i_ determinations for each batch of *ee*AChE in an effort to avoid a substantial change of the specific enzyme activity during the entire course of multiple *K*_i_ determinations. Second, because nonspecific binding of a test inhibitor to microsomes, phospholipid, and albumin can affect the *IC*_50_ and *K*_i_ of the inhibitor [28–30], we excluded albumin, glycerol, gelatin, or any other enzyme stabilizers in all of our enzyme inhibition assays reported in this article to minimize nonspecific binding of a test inhibitor. Third, according to our adsorption studies above, we also excluded the use of glass containers for inhibitor stock solutions to avoid the adsorption-caused deviation of the actual inhibitor concentration from the nominal inhibitor concentration. Reassuringly, Table 2 shows that the increase of *K*_i_ of tacrine or bis(7)-tacrine is indeed inversely proportional to the increase of the specific enzyme activity at the time when the assay was performed. This inverse correlation shows that inactive enzyme can serve as a nontarget binding site for a test inhibitor during the process of the enzyme inhibition assay. It explains that the *K*_i_ variations for both tacrine and bis(7)-tacrine were due to the reduction of the actual inhibitor concentration caused by the inhibitor binding to the inactive enzyme relative to the nominal inhibitor concentration. It also suggests that the reported *K*_i_ variations for tacrine [15–18] were caused likely by the inhibitor binding to the inactive enzyme.

**Table 2.** Effect of specific enzyme activity on *K*_i_

| *eeAChE*<sup>a</sup> in pg | SEA<sup>b</sup> in U/mg (mean ± SE<sup>c</sup>) | $K_i$<sup>d</sup> (mean ± SE<sup>c</sup>) |
| --- | --- | --- |
| Bis(7)-tacrine |  |  |
| 10.4 | 342 ± 10 | 2.9 ± 0.4 pM |
| 41.7 | 252 ± 6 | 81 ± 2 pM |
| 125.0 | 45 ± 2 | 457 ± 23 pM |
| 250.0 | 26 ± 2 | 734 ± 70 pM |
| Tacrine |  |  |
| 10.4 | 342 ± 7 | 11 ± 1 nM |
| 41.7 | 225 ± 23 | 14.6 ± 0.3 nM |
| 83.3 | 124 ± 7 | 32 ± 1 nM |
| 125.0 | 46 ± 7 | 44.6 ± 0.8 nM |
| 250.0 | 27 ± 3 | 72 ± 5 nM |

<sup>a</sup>*eeAChE*: the amount of *Electrophorus electricus* acetylcholinesterase in the assay solution.
<sup>b</sup>SEA: specific enzyme activity of *EeAChE* that was determined using the data in Table S1 at pH = 8.0, 50 mM sodium phosphate buffer, temperature of 26 ± 2 °C, and acetylthiocholine concentration of 2.0 mM. <sup>c</sup>SE: standard error of five independent experiments. <sup>d</sup> $K_i$ was determined from the data in Table S1 and Fig. S1 using plastic vials for inhibitor stock solutions.

### 3.3. Caveat of using the experimental K_i_ as an absolute affinity indicator

The above studies offer the first set of experimental evidence for container surfaces and inactive enzymes to serve as nontarget binding sites for test inhibitors. These studies demonstrate the profound but long-overlooked effects of these nontarget sites on the experimentally determined *K*_i_. Because the binding of a test inhibitor to the container surface during the inhibitor solution preparation process and/or to the inactive enzyme during the enzyme inhibition assay process is not factored in conventional experimental determination of *K*_i_, we caution against using the experimentally determined *K*_i_ as a measure of absolute binding affinity to rank order binding potencies, define selectivity, or benchmark computational methods without knowing detailed assay conditions. To facilitate the use of the experimentally determined *K*_i_ as an “absolute” binding affinity indicator, we suggest greater evaluation and optimization of assay conditions to minimize inhibitor binding to nontarget sites as well as reporting, in online supplementary documents, all experimental details including specific enzyme activities at times when assays are performed (see Ref. [31] for an excellent example of reporting specific AChE activity associated with the reported *K*_i_ of tacrine).

## Conflict of interest

The listed authors have no conflict of interests.

## Author contributions

Y.-P.P. conceived, designed, and supervised the project. Y.-P.P. and F.H.D. designed the UV absorption protocol and revised the enzyme inhibition assay protocol described in Ref. [5]. Y.-P.P. synthesized bis(7)-tacrine. F.H.D. performed all enzyme inhibition and UV absorption assays, observed the correlation between *K*_i_ and the amount of *ee*AChE required for the *K*_i_ determination, and drafted the methods section. Y.-P.P. and F.H.D. analyzed and interpreted all experimental data. Y.-P.P. wrote the paper. Y.-P.P. and F.H.D. contributed with revisions.

## Acknowledgments

This work was supported by the US Army Research Office (W911NF-16-1-0264) and the Mayo Foundation for Medical Education and Research. The contents of this article are the sole responsibility of the authors and do not necessarily represent the official views of the funders.

